# A rapid, automation-friendly, cell-free and purification-free pipeline for binding and kinetic characterisation of de novo designed binders with high quality standardised per-design data

**DOI:** 10.64898/2026.10.19.759856

**Authors:** Diego Ruiz Espi, Gustavo Avelar Molina, Oliver Sonne Buch, Julia Heyd, Aske Unger, Line Winther Gormsen, Katrine Boldsen Lund, Kritika Sahni Ray, Charles Williams, Andreas Hald, Timothy P. Jenkins, Esperanza Rivera-de-Torre

## Abstract

De novo protein binders can now be generated far faster than they can be tested. Current validation routes — display selection, cell-free two-hybrid assays, or surface-based biophysics — require cell culture, colony picking, protein purification, target labeling or sensor immobilisation, take 2 to 8 weeks, and require additional methods for affinity and kinetics characterisation. Here we report an end-to-end, cell- and purification-free pipeline that goes from gene to binding affinity and kinetics in three days without genetically modified organisms, cell culture, purification, labelling or immobilisation. An automated, metric-driven loop of RFdiffusion, ProteinMPNN and AlphaFold2 initial-guess generates and filters designs, which are ranked by Boltz-2 ipSAE, assembled by Golden Gate cloning from linear DNA, expressed in a cell-free protein synthesis lysate as fluorescent protein fusions, and screened directly in the unpurified product by flow-induced dispersion analysis (FIDA). Against human growth hormone, 11 of 30 designs (37%) bound, and the four strongest had equilibrium dissociation constants of 0.93–9.98 nM with on- and off-rates resolved from the same cell-free material. Each sample run measures expression level, hydrodynamic radius, oligomeric and aggregation state alongside the binding readout, producing the standardised, per-design datasets that model training requires.

## Introduction

Proteins are now designed faster than they can be tested. Generative structure and sequence design tools [1–5] have brought protein binder design within reach of any group with a GPU [6]. Yet, even after filtering with several in silico confidence metrics [7–13], reported success rates have ranged from 7.5% in the first round of the Adaptyv EGFR competition [14] to a mean of 46.3% across 12 targets for BindCraft [5]. This means both that in vitro validation is still a must, and that a single 96-well plate would be expected to contain at least 2 binders based on these success rates. The bottleneck is therefore not the number of designs that can be tested but the format of the assay: its throughput, workload and data quality.

Different binder screening methods provide solutions to different needs. Display methods (phage, yeast, cDNA and mRNA) can test thousands to tens of thousands of designs in a single run. A typical display-based validation run requires 2 to 8 weeks and comprises many labour-intensive steps that can fail i.e. cell transformation, cell culture, colony picking, protein purification, and deep sequencing [15–20]. Each step brings its own attrition rate, lowering the fidelity of designs to the final dataset, finally only selecting tens of designs for affinity and kinetic characterisation, requiring an additional round of transformations, protein expression and purification for analysis by e.g., surface plasmon resonance (SPR) or biolayer interferometry (BLI). Display methods are thus ideal for testing thousands to millions of designs where the expected success rate is extraordinarily low, but are excessive in most cases for the current paradigm of computation-led design campaigns. Moreover, display methods value throughput over data quality, which reduces learnings from the model-training perspective [21–24]. They only yield a binary yes/no binding data point per binder design, which heavily depends on the concentration of binding target used. In addition, it is not a straightforward task to deconvolute non- or low-expression from non-binding.

Recent efforts to bridge this gap include the cell-free two-hybrid assay (CF2H) to screen designs without cloning or purification, however, the target must be biotinylated and fused to a DNA-binding domain, requiring additional target expression, purification and biotinylation labeling steps. Binding readouts are then retrieved from a fold-change in fluorescence that does not account for binder expression level. Finally, the approach relies on additional methods for validating binder expression, affinities and kinetics, again involving more expression and purification steps [25].

An approach to avoid purification steps is to apply flow-induced dispersion analysis (FIDA) [26] to analyze heat-treated bacterial cell lysates for identification of binders in-solution without immobilisation, avoiding sensor-surface artifacts. However, there remain the limitations that each target must be fluorescently labeled, the heat step selects simultaneously for expression, binding, and thermal stability, which are difficult to deconvolute and affinities are still obtained only after re-expression, purification and analysis through other biophysical techniques [27].

In summary, current methods for binder screening and affinity characterisation rely on cell-based expression, transformation, colony picking and protein purification, which are slow and labour-intensive over a 2 to 8 week timeline. The protein design field lacks an in vitro validation pipeline that is fast, automation-friendly, inexpensive and simple enough to set up in a standard molecular biology laboratory, without genetically modified organisms or specialised display-selection expertise. The pipeline should additionally return standardised data valuable for model training. To achieve this, the assay should work across designs with little optimisation, modification or preparation, and should require minimal data treatment, and in which the oligomeric state, and aggregation tendency of each partner is defined and consistent across designs.

Here we report an automatable, cell- and purification-free pipeline that strives to meet these requirements (Fig. 1). Designed sequences are assembled by Golden Gate cloning from linear DNA, expressed in a cell-free lysate [28] and screened by FIDA directly with the unpurified product to yield the initial verification of binding. Best-performing candidates are then re-analysed for the affinity and on- and off-rates from the same cell-free protein synthesis (CFPS) material, all within three days from receiving DNA and without genetically modified organisms, cell culture, purification, immobilisation or extra labeling steps. Binder expression, size, oligomeric state and aggregation state are validated and quantified, all in the same experiment, providing a richer dataset of both binder characteristics and binding affinity data ideal for model training [21–24]. To test this pipeline, a target structure and a GPU, an automated loop of RFdiffusion [1], ProteinMPNN [2] and AlphaFold2 initial guess (AF2-IG) [3] generates and filters binders, which are then ranked with Boltz-2 ipSAE [11, 12]. As the designs are chosen using in silico confidence metrics, we also ask whether those metrics still rank the designs that pass them. We validated the pipeline on human growth hormone (hGH), a well-characterised system whose Site-1 epitope was mapped three decades ago [29, 30] and is already exploited therapeutically by the growth hormone receptor (GHR) antagonist pegvisomant [31]. Of 32 designs, 30 were successfully expressed and 11 (34%) bound, and an unseeded design campaign produced more passing designs in less time than hot-spot-seeded campaigns.

**Fig. 1.**
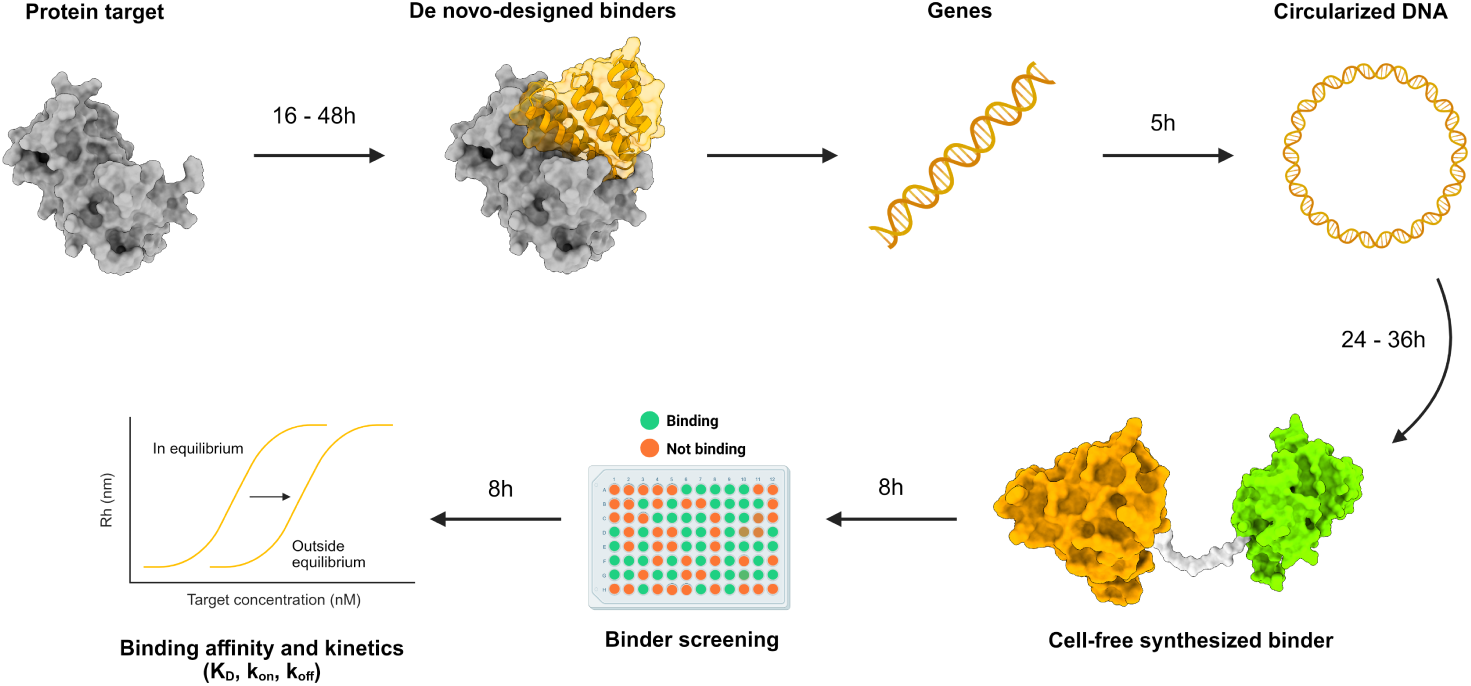
The automatable, iterative, metric-driven optimisation pipeline designs binders with the desired in silico metrics against any target of interest. The top ranked binders are ordered as genes (dsDNA) and then circularised with the cell-free protein synthesis (CFPS) expression vector. Binders are then expressed fused to mChartreuse and screened for binding against their target by flow-induced dispersion analysis (FIDA). Finally, the most promising candidates undergo binding affinity and kinetics characterisation using the same cell-free material.

## Results

### Automated iterative metric-driven optimisation for binder designs successfully generates binders with the requested in silico metrics

The overarching objective was to design and validate a pipeline end-to-end, from automated iterative metric-driven binder design and optimisation, through to binder screening and characterisation. We first benchmarked the design stage across three campaigns, then used the selected designs to establish CFPS and FIDA screening, and finally asked whether any in silico metric could have predicted which designs would bind. We ran three protein design campaigns: one blind run (unseeded) and two hot-spot rounds, HS 1 (His21, Phe25, Arg167) and HS 2 (Leu6, Leu9, Thr123). All three were given the same objective: design at least 100 binders with ipAE *<* 8 Å and pLDDT *>* 80 within a maximum of five partial diffusion rounds.

Each campaign was driven by the same automated loop (Fig. 2a): RFdiffusion generated 300 candidate scaffolds against the target interface (either unseeded or with the selected hot-spots), ProteinMPNN generated 5 sequences per scaffold, and the folding of each pair of sequences was predicted and scored with AlphaFold2 Initial Guess (AF2-IG). Designs that met ipAE *<* 8 Å and pLDDT (AF2-IG) *>* 80 (Fig. 2b, top) were retained; the top 15 (maximum 2 per scaffold) were used as starting point for a subsequent round of RFdiffusion partial diffusion. This scaffold-sequence-fold filter cycle was repeated for up to 5 rounds per campaign until at least 100 designs had passed the ipAE/pLDDT thresholds. The resulting designs were then confirmed and re-evaluated with a second, independent structure-prediction in silico metric (Boltz-2 ipSAE min) as a final filter before proceeding to in vitro validation (Fig. 2a).

**Fig. 2.**
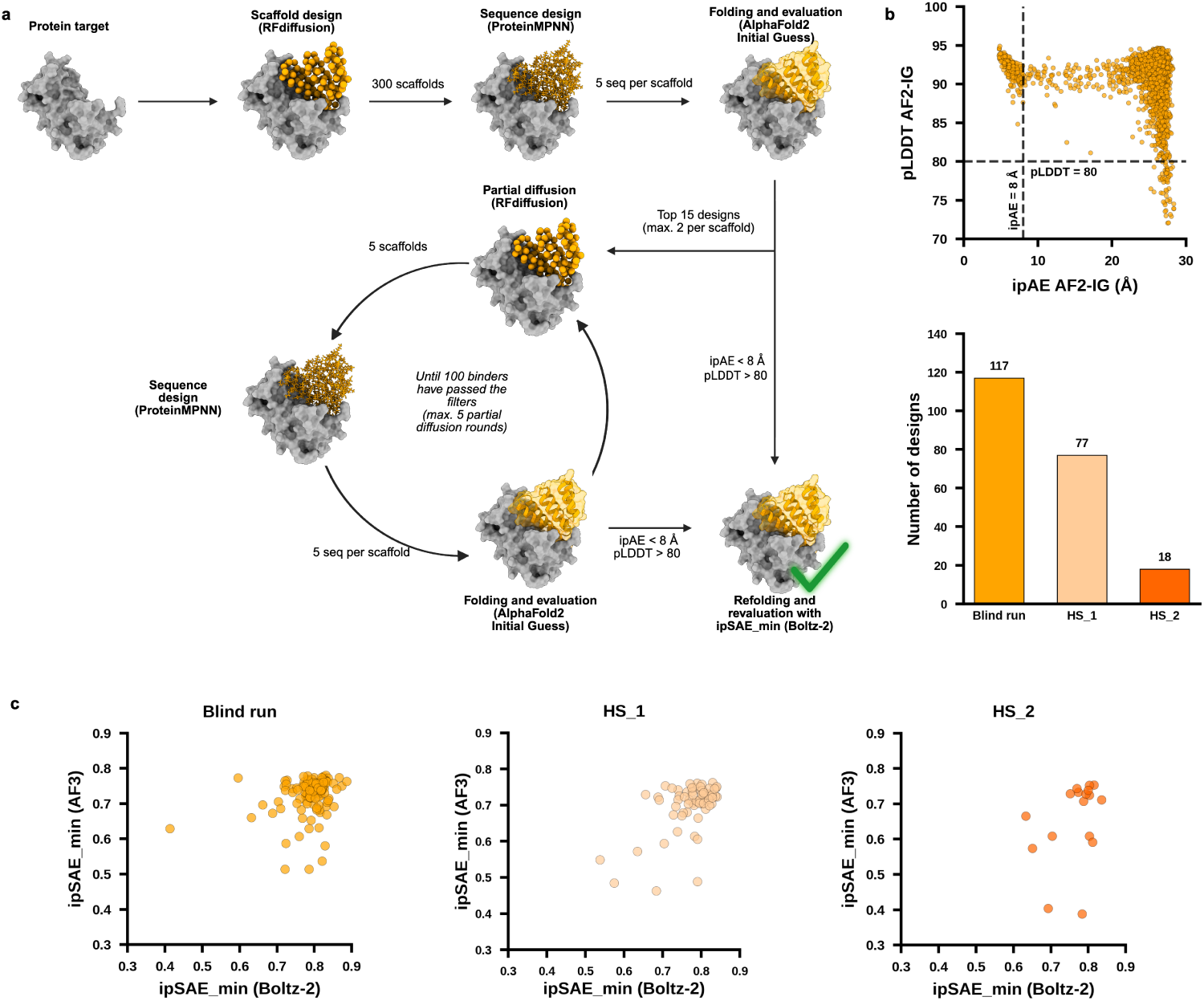
**a,** The in silico pipeline starts with a given PDB file with the structure of the protein target, and optionally, the in wished in silico metrics, and the number of scaffolds and sequences per scaffold in each round. Once the run is started, RFdiffusion generates scaffolds, which are then taken by ProteinMPNN to generate a number of sequences per scaffold, which fold will be predicted and evaluated based on its ipAE and pLDDT scores by AF2-IG. Then, the top candidates are partially diffused (RFdiffusion) and a number of sequences are generated for each new diffused scaffold, which fold is predicted and evaluated with AF2-IG, and the top candidates go for another partial diffusion round. These partial diffusion rounds go on until the number of designs within the set thresholds is equal or higher than the defined, or if the maximum number of partial diffusion rounds is reached. Once the partial diffusion rounds are done, Boltz-2 revaluates the designs and ranks them by ipSAE min. **b,**Top panel, the ipAE and pLDDT of all the binders generated in all three design campaigns. Bottom panel, the number of designs per campaign that passed the wished filters (ipAE *<* 8 Å and pLDDT *>* 80). **c,** Designs passing the AF2-IG thresholds revaluated with Boltz-2 and AlphaFold3 ipSAE min for each of the design campaigns.

The blind run stopped after the first partial diffusion round, having generated 375 scaffolds and 1,875 sequences, of which 117 unique-scaffold designs passed the requested filters. HS 1 and HS 2 ran all 5 partial diffusion rounds, generating 675 scaffolds and 3,375 sequences each, of which 77 and 18 designs passed the requested filters in HS 1 and HS 2, respectively (Fig. 2b). The blind run finished after 16 hours, whereas the hot-spot campaigns ran for 36 hours on an NVIDIA A100 Tensor Core GPU. The blind run therefore generated 52% and 550% more designs than HS 1 and HS 2, respectively, in just 44% of the time.

The designs from the three campaigns performed similarly when evaluated with ipSAE min Boltz-2 and AlphaFold3 (AF3), with most designs across all three campaigns scoring between 0.7 and 0.8 on both metrics (Fig. 2c). Across the pooled 201 designs, Boltz-2 gave a slightly higher mean ipSAE min than AF3 in every campaign (pooled: 0.781 vs 0.710, gap 0.072) (Fig. 2c). Computed separately within each campaign (Fig. 2c), the two metrics were positively correlated in the blind run (Pearson *r* = 0.24, *R*^2^ = 0.06, *p* = 8.9 *×* 10*^−^*^3^; Spearman *ρ* = 0.23, *p* = 1.5 *×* 10*^−^*^2^; *n* = 114) and, more strongly, in HS 1 (Pearson *r* = 0.57, *R*^2^ = 0.33, *p* = 2.3 *×* 10*^−^*^7^; Spearman *ρ* = 0.35, *p* = 3.2 *×* 10*^−^*^3^; *n* = 70); in HS 2 the correlation did not reach significance (Pearson *r* = 0.37, *R*^2^ = 0.14, *p* = 0.14; Spearman *ρ* = 0.31, *p* = 0.22; *n* = 17), consistent with that campaign’s small size. Pooled across all three campaigns, the two metrics remained positively correlated (Pearson *r* = 0.39, *R*^2^ = 0.15, *p* = 1.1 *×* 10*^−^*^8^; Spearman *ρ* = 0.30, *p* = 1.8 *×* 10*^−^*^5^; *n* = 201).

We ranked the binders by Boltz-2 ipSAE min and selected the top 32 (four full columns of a 96-well plate) for in vitro validation. Of these, 20 came from the blind run, 9 from HS 1 and 3 from HS 2, corresponding to 17%, 12% and 17% of the designs that passed the initial thresholds, respectively.

### Rapid, low-volume cell-free protein-synthesis from circularised DNA yields high, quantifiable binder expression levels

With the 32 top-ranked designs in hand, we developed a rapid, readily automatable GMO-free molecular cloning and expression pipeline (Fig. 3a). The binder genes were reverse-translated, codon-optimised for *Nicotiana tabacum* and ordered as dsDNA gene fragments. Genes were amplified by PCR with consistent yields of 80-120 ng/µL after purification. Subsequent Golden Gate assembly similarly yielded purified DNA concentrations of 72-120 ng/µL, well above the 50 ng/µL recommended by the supplier of the ALiCE CFPS (LenioBio). These assembly products comprised a range of vector (V)-gene (G) assemblies (e.g. V-G, G-V-G, V-G-V-G, etc.), all of which are valid DNA templates for expressing binders (Fig. 3c).

**Fig. 3.**
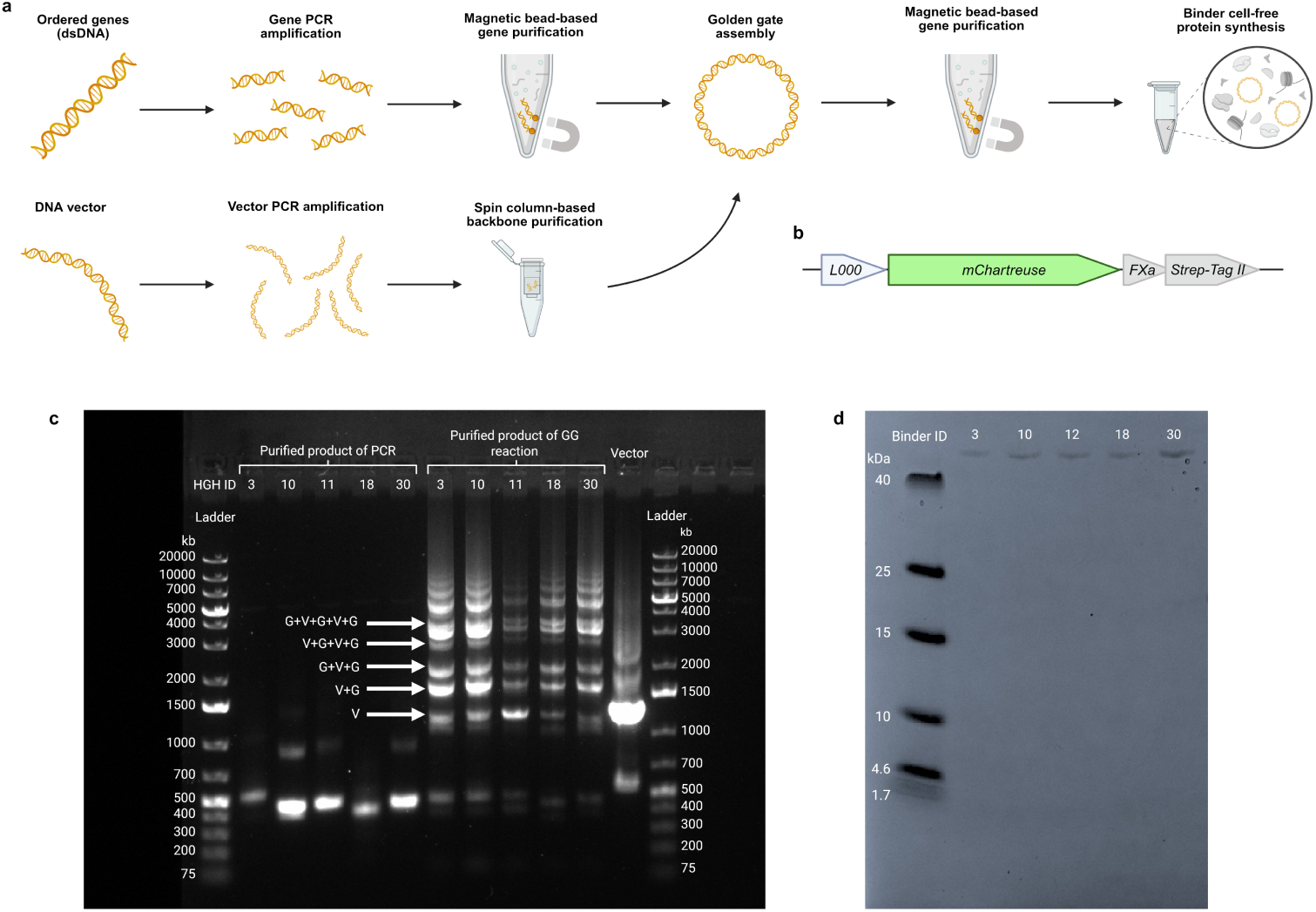
**a,** The dsDNA gene fragments coding for the binders are amplified by PCR and purified with magnetic beads. Amplified genes are circularised by Golden Gate (GG) assembly with the vector containing essential transcription and translation regulatory elements, mChartreuse and Strep-Tag II in phase with the open reading frame. A final magnetic bead purification, the construct is produced in cell-free expression. **b,** Linear representation of the part of the vector that is expressed, detailing the linker, mChartreuse, an FXa protease cleavage site and the Strep-tag II for protein purification. **c,** Agarose gel electrophoresis result of the purified PCR and GG reaction products for 5 of the binders (HGH 3, HGH 10, HGH 11, HGH 18 and HGH 30) together with the purified vector. The purified PCR product bands correspond to the correct sizes of the binders’ DNA (450-550bp), whereas the vector brightest band also corresponds to the correct length (1,269bp). In the purified GG products, we can see that most of the DNA has been circularised into all the different possible species: vector+gene (V+G), gene+vector+gene (G+V+G), vector+gene+vector+gene (V+G+V+G), etc. **d,** SDS gel of the same expressed and purified binders (expected size 42.1 kDa).

We expressed the binders as C-terminal fusions to mChartreuse and Strep-tag II using the CFPS system (Fig. 3b). Five binder candidates were purified by StrepTactin magnetic beads and analysed by SDS-PAGE gel to confirm their size in an orthogonal method and compare it to FIDA measurements. To measure expression yields without purifying every binder, we analysed one of the purified candidates by FIDA at a range of concentrations to build a calibration curve of the area under the curve (AUC), showing a linear correlation of *R*^2^=0.997. The measured concentrations of the unpurified binder panel ranged from 7.6 to 121.9 nM (760 nM to 12.19 *µ*M before the 100-fold dilution) (Fig. 4d), whereby 2.5 nM was identified as the minimum concentration required to maintain a signal-to-noise ratio above 100. This quantitative calibration (Fig. 4e) was essential for all subsequent binding-affinity and kinetics experiments, ensuring that the measured K*_D_* and rate constant values reflect true binding properties rather than concentration artefacts. The commercially sourced target protein (hGH) also underwent a fast quality control via FIDA, with 280nm detector, and showing the expected hydrodynamic radius (2.4 nm), and only a few aggregated species detected as events between minute 0.9 and 1.3 (Fig. 4h). For this, we used the same FIDA method as the baselines in the binding screening.

**Fig. 4.**
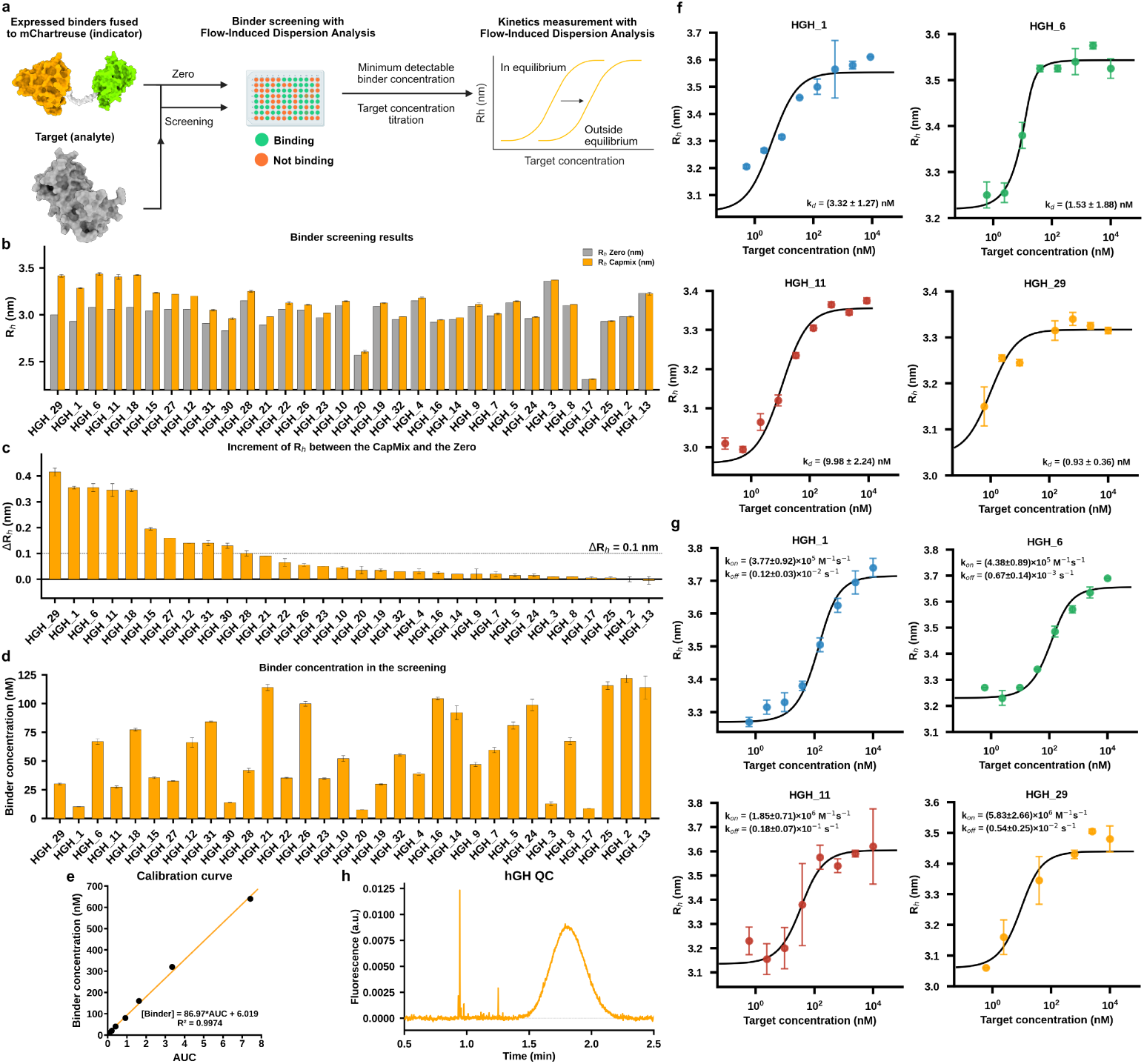
**a,** The binders were analysed by FIDA with a 1:100 dilution of the CFPS material, with and without the target analyte hGH. An increase in hydrodynamic radius is expected when the target analyte is present in the case of binding. Then, the top candidates are run twice more at minimum detectable binder concentration against a titration of the target analyte: once in PreMix mode to measure the binding affinity in equilibrium, and once with high-pressure CapMix mode to analyse the binding before it reaches the equilibrium and measure kinetics. **b,** Binder screening results: hydrodynamic radius measured for each binder with and without the target analyte. **c,** Increment of hydrodynamic radius between the zero (binder alone) and the CapMix (binder + target analyte). **d,** Binder concentration in the screening measured with the calibration curve based on fluorescence AUC). **e,** Calibration curve constructed from a purified binder-mChartreuse fusion protein to enable concentration measurements in unpurified samples. **f,** Binding curves in equilibrium (PreMix) of the top 4 binders showing the highest increment on hydrodynamic radius between the zero and the CapMix mode, with the binding affinity measurements. **g,** Binding curve outside equilibrium of the top four binders showing the highest increment on hydrodynamic radius between the zero and the CapMix mode, with the binding kinetics measurements. **h,** Target Quality Control: hGH run with the 280 nm detector to verify its hydrodynamic radius (2.4 nm), oligomeric state (monomer) and potential aggregates (at least two species).

### FIDA allows for rapid, low-volume screening of unpurified CFPS material and resolves binding affinity and kinetics from the same sample

We subsequently aimed to develop a clean, readily automatable method for binder screening that requires no purification (Fig. 4a). We diluted the CFPS product 1:100 and used a volume of 20 µL to avoid potential capillary clogging by CFPS-derived host-cell proteins or aggregates and used FIDA in CapMix mode to measure the hydrodynamic radius of the binder-mChartreuse fusion proteins alone and in the presence of a constant concentration of hGH (5 µM), both in duplicate. The first measurement confirmed successful expression and a suitable hydrodynamic radius (*R_h_*) of 2.9-3.1 nm for 30 of the 32 binder fusion proteins (Fig. 4b). It also allowed us to calculate the concentration of each expressed fusion protein from the fluorescence AUC and the calibration curve, again without any purification, giving concentrations between 760 nM and 12.19 *µ*M (Fig. 4d and e). By contrast, HGH 17 and HGH 20 had hydrodynamic radii smaller than the 3 nm expected for the intact binder–mChartreuse fusion. HGH 17 showed an *R_h_* of 2.3 nm, matching free mChartreuse, indicating that the binder had been completely cleaved from the fusion; HGH 20 showed an *R_h_* of 2.6 nm, intermediate between free mChartreuse and the expected fusion size, consistent with partial cleavage. Because the expected fusion size could not be confirmed for either design, both were excluded from the binding study.

We then calculated the increase in the hydrodynamic radius (Δ*R_h_*) between the two measurements with and without the hGH target and found that 11 of the 30 binders showed an increase in the hydrodynamic radius greater than 0.1 nm, ranging from 0.43-0.11 nm (Fig. 4c). The in vitro binding success rate was therefore 37%, and 94% of binders expressed correctly.

To determine binding affinities and kinetics, we performed two further rounds of FIDA on the four binders with the largest Δ*R_h_* (HGH 1, HGH 6, HGH 11, and HGH 29), using the same CFPS material diluted to a fixed binder concentration against an 8-point titration of hGH (0.61 nM - 10 µM). The first round used Premix mode to obtain an equilibrium binding curve; binder and target were mixed in advance so that we could measure the hydrodynamic radius of the complex could be measured at equilibrium. The four binders showed low-nanomolar affinity, with K*_D_* ranging from 0.93 ± 0.36 nM to 9.98 ± 2.24 nM (Fig. 4f). The second round used CapMix with increased injection pressure, moving the binding reaction out of equilibrium, here the k*_on_* and k*_off_* constants showed were more diverse. k*_on_* ranged from 3.77 ± 0.92 × 10^5^ M*^−^*^1^s*^−^*^1^ to 5.83 ± 2.66 × 10^6^ M*^−^*^1^s*^−^*^1^, while the k*_off_* ranged from 0.67 ± 0.14 × 10*^−^*^3^ s*^−^*^1^ to 0.18 ± 0.07 × 10*^−^*^1^ s*^−^*^1^ (Fig. 4g).

### In silico confidence metrics saturate after filtering with AF2-IG ipAE and pLDDT, and Boltz-2 ipSAE min, and no longer rank experimental binders

Finally, to assess whether an additional in silico filter could have increased in vitro success, we ran a small meta-analysis of the 32 tested binders. Campaign identity (hot-spot vs blind run) did not predict in vitro success (8/20, 2/9 and 1/3 for the blind, HS 1 and HS 2 campaigns, respectively; P = 0.65, Fig. 5e), nor did the specific backbone trajectory: the hit rate ranged from 0% to 100% across the 11 trajectories, but this spread was statistically indistinguishable from binomial sampling noise at the group sizes tested (permutation test, P = 0.40) (Fig. 5a). The designs also converged structurally and physicochemically, regardless of outcome. Every trajectory, including those from the blind campaign, converged on the same face of hGH, contacting a common set of three surface patches, with four residues — Gln18, Gln22, Asp25 and Asp26 — engaged by all 32 designs irrespective of outcome (Fig. 5b); no individual residue’s contact frequency discriminated binders from non-binders after correction for multiple comparisons (q 0.41 across 68 positions, Fig. 5h). This convergence extended to the physicochemical basis of the interaction: both binders and non-binders relied on the same electrostatic strategy, dominated by binder acidic residues pairing with hGH arginines (mean 2.91 versus 1.86 Glu–Arg salt bridges per design, Fig. 5g), and no hGH residue reached significance as a binder-specific polar or electrostatic anchor after correction (all q = 0.72). It is worth to mention that the pre-synthesis metrics themselves had no residual predictive power. Neither static interface geometry (Fig. 5c) — buried interface area, hydrogen-bond count and helical content were indistinguishable between groups (P = 0.66, 0.51 and 0.03, respectively, the last in the direction opposite to expectation)— nor pre-synthesis structure-prediction confidence (Fig. 5d) anticipated success: AF2-IG ipAE, Boltz-2 ipSAE min and AlphaFold3 ipSAE min all failed to separate outcomes (P = 0.50, 0.55 and 1.00; area under the receiver-operating characteristic curve 0.58, 0.43 and 0.50, respectively), with every design — successful or not — occupying a single narrow, high-confidence region of prediction space. This ceiling effect is consistent with the fact that these metrics had already been used to filter the candidate pool before synthesis (Fig. 5f).

**Fig. 5.**
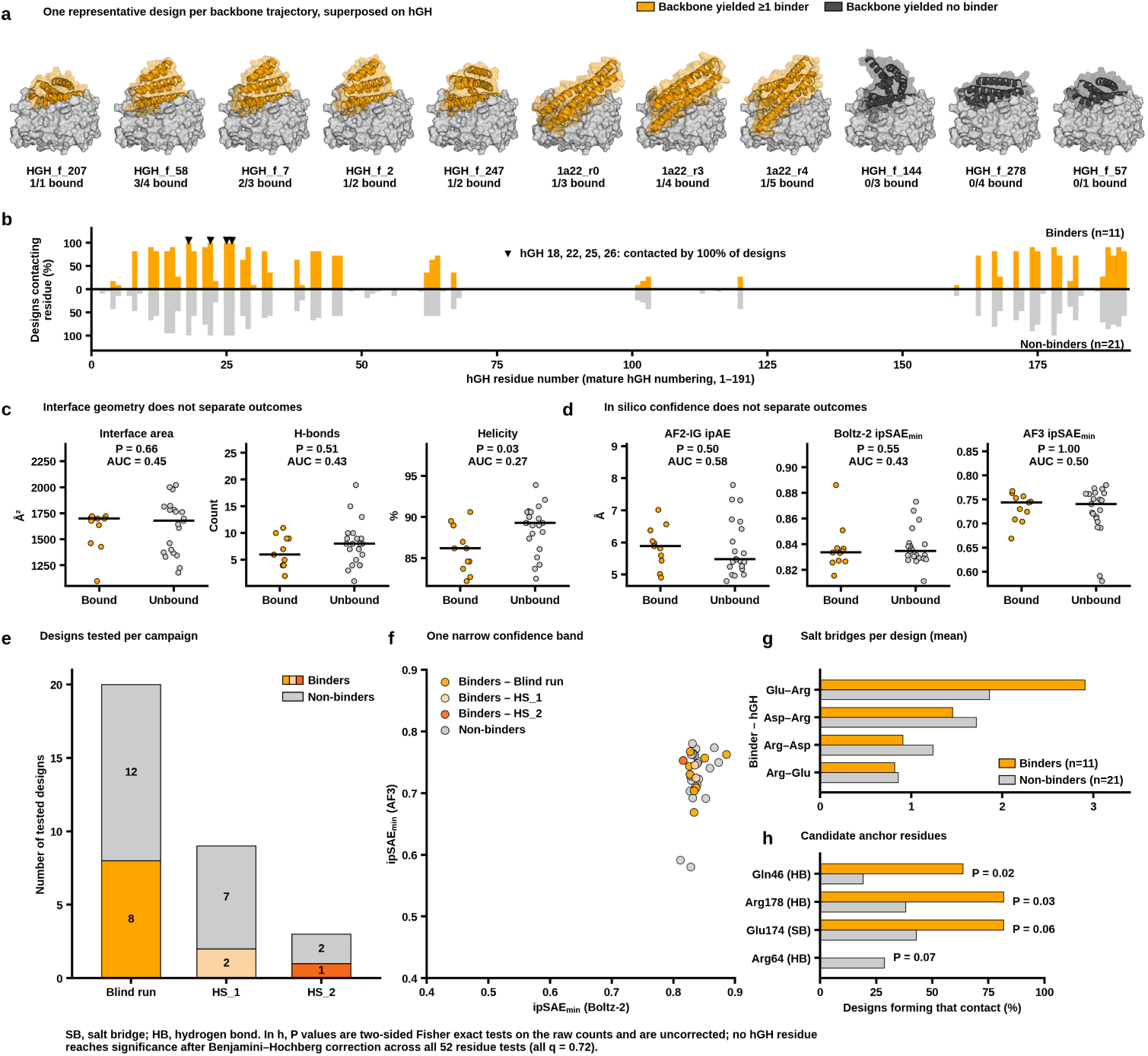
**a**, One representative design per RFdiffusion backbone trajectory (*n* = 11), superposed on hGH; orange, trajectory yielded one or more than one confirmed binder; grey, yielded none. Fraction bound given per trajectory. **b**, Per-residue hGH contact frequency for binders (orange, *n* = 11) and non-binders (grey, *n* = 21); arrowheads mark residues 18, 22, 25, 26 (contacted by 100% of designs). **c**, Interface area, H-bond count and helicity, bound vs. unbound (two-sided Mann–Whitney *U*; AUC, classifier performance). **d**, Pre-synthesis confidence metrics (AF2-IG ipAE, Boltz-2 ipSAE min, AF3 ipSAE min), bound vs. unbound. **e**, Designs tested per campaign (blind run, HS 1, HS 2); hit rates 8/20, 2/9, 1/3 (Fisher’s exact *P* = 0.65). **f**, AF3 vs. Boltz-2 ipSAE min, all 32 tested designs (Pearson *r* = 0.33, *P* = 0.06; Spearman = 0.17, *P* = 0.35). **g**, Mean salt bridges per design for the four most frequent binder–hGH residue pairs, bound vs. unbound. **h**, The four hGH residues most differentially engaged by binders versus non-binders in a specific contact: Gln46 and Arg178 by hydrogen bond and Glu174 by salt bridge, all enriched in binders, and Arg64 by hydrogen bond, seen only in non-binders (two-sided Fisher’s exact, uncorrected; none survives Benjamini–Hochberg [32] correction across the 52 residues tested, all *q* = 0.72). SB, salt bridge; HB, hydrogen bond.

## Discussion

Here, we developed and validated an end-to-end pipeline that meets the needs of the current protein binder design field: a molecular biology laboratory with access to a flow-induced dispersion analysis (FIDA) instrument can now generate and validate its own de novo binder designs with standard bench equipment, starting from nothing more than a target structure. The in silico stage is fully automated - given a PDB file and a single GPU, an iterative RFdiffusion-ProteinMPNN-AF2-IG loop runs until the ipAE and pLDDT objectives are met, and Boltz-2 ipSAE min then serves as an independent final filter, with no coding required. The in vitro, cell-free validation stage takes three days from gene to binding kinetics with minimal hands-on work and uses no genetically modified organisms, extra labelling steps, cell culture or protein purification. The entire pipeline is automatable but also provides timesaving advantages for manual handling. With a single FIDA instrument, three 96-well plates (288 designs) can be processed in parallel every 5 days, as new plates can be initiated once the previous plate has completed the first PCR step. Golden Gate templates assembled from linear DNA are expressed in a CFPS lysate and screened directly, with FIDA supplying initial binding confirmation for every design and subsequent equilibrium affinity and on-and off-rates for those best-performing binder candidates selected for follow-up, all from the same unpurified CFPS material and under identical conditions. Against human growth hormone (hGH), 30 of the 32 binder candidates were successfully expressed and 11 designs successfully bound the target (37%). The four binders with the largest radius shifts had *K_D_* values of 0.93–9.98 nM. Considering that each design is expressed, quantified, size-checked and measured in the same way, the failures are as informative as the successes, yielding the standardised, per-design data that model training requires.

The design campaigns indicated that seeding with hot-spot was unnecessary for this target. Every design that passed the initial filters landed on the same face of hGH, the receptor-binding face [29, 30], whether or not the campaign had been told where to bind. The unseeded run nonetheless produced many more designs in less time: it met the 100-design objective after a single partial-diffusion round, returning 117 unique-scaffold designs in 16 h, whereas the two hot-spot campaigns used all five rounds and 36 h on the same GPU to return 77 and 18 designs. The blind run therefore yielded 52% more passing designs than HS 1 and 6.5 times as many as HS 2, in 44% of the compute time, and experimental success did not differ between campaigns (8/20, 2/9 and 1/3; *P* = 0.65). A plausible interpretation is that, for a dominant solvent-exposed epitope, the design network already favours the productive face, so that hot-spot constraints mainly shrink the set of scaffolds able to satisfy the filters.

Compared with the screening approaches previously described for testing designed proteins, the method described herein reduces the broad scales achievable by display methods for speed, simplicity and higher data quantity and quality per design. Display technologies yield just single, yes/no binary binding data points from many binders after 2-8 weeks of intensive workload, which is useful when you are expecting an extremely low success rate in your campaign, but not as useful for generating high quality data sets, as many questions such as binder expression and the definition of binding are not standardised, and affinity and kinetics are not measured [17, 18].

The CF2H method reduces workload but still requires colony picking, target protein expression and purification. It does not provide a solution for verifying binder expression, and in some cases, target aggregation through its fusion partner ends in unconclusive results [25].

Closest to our method is FIDA of heat-treated bacterial cell lysates. The readout principle is shared but we designed our pipeline to avoid its limitations. The published FIDA screening method requires each protein target to be fluorescently labelled as the indicator, whereas we express the binder fused to the label and use unlabelled target, eliminating the additional labeling step. Screening of cell lysates does not normalise to expression, and a low-µM binder was missed for this reason, whereas the fluorescence of our binder fusion quantifies expression in the same measurement. The heat treatment step also retains only thermostable designs and the method requires genetically modified *E. coli* cell culture for cloning and recombinant protein production, whereas our templates are assembled and expressed from linear dsDNA without transformation or living cells. The shared trade-off of using FIDA as the readout with fluorescent binders as the indicator is that binding is read as a change in hydrodynamic radius, which can be challenging when the expected increment in hydrodynamic radius is lower than 5% in nanometers. [27].

None of these published methods are able to measure binding affinity nor kinetics [17, 18, 25, 27]. Typically, they require an additional method such as BLI or SPR, where each design, requires expression, purification and immobilisation on a sensor, thus limiting panels to tens of designs and adding weeks between initial screening and affinity and kinetic measurements. In addition, BLI and SPR do not report aggregation, and oligomerization can be difficult to determine [21] - BLI in particular can report false positives of non-specific binding as on-target binding. Our method measures binding affinity and kinetics from the same material, and reports on temperature, aggregation, size/oligomerisation and binder concentration in every single measurement. Additionally, measurement of unpurified CFPS material also provides an internal check on non-specific binding. All these parameters contribute to a method best suited for building high-quality data sets for model training to date.

Across all 30 successfully expressed designs, the binder-mChartreuse fusion proteins on their own had hydrodynamic radii of 2.9–3.1 nm, matching the theoretical size of a globular protein of their length; none associated with lysate proteins strongly enough to shift the radius, and a radius increase (0.43–1.11 nm in the 11 binders) appeared only on adding hGH, against the background of the full lysate. None of the designs formed large aggregates. This is a screen for gross non-specific interaction rather than a substitute for testing against a panel of unrelated targets, but purified-protein formats do not provide even this, and it is an informative readout given the amount and variety of molecules present in the lysate (including proteins, nucleic acids, free nucleotides, and amino acids). The same material and technique was used to resolve kinetic diversity: among the four strongest binders, *K_D_* spanned about 10-fold (0.93 ± 0.36 nM to 9.98 ± 2.24 nM), whereas *k_on_* varied about 15-fold (3.77 ± 0.92 × 10^5^ M*^−^*^1^s*^−^*^1^ to 5.83 ± 2.66 × 10^6^ M*^−^*^1^s*^−^*^1^) and *k_off_* about 27-fold (0.67 ± 0.14 × 10*^−^*^3^ s*^−^*^1^ to 0.18 ± 0.07 × 10*^−^*^1^ s*^−^*^1^), even though binders and non-binders alike are predicted to use the same acidic-residue-to-arginine electrostatic strategy on the same epitope.

The value of these per-design measurements lies in what they permit one to conclude from a negative result. Published affinity values frequently omit the controls and conditions needed to interpret them [22, 24], and apparent affinity is confounded by the binding-competent fraction and by oligomeric state, which together can shift *K_D_* and *k_off_* by an order of magnitude [21, 24]. Trained on such data, models learn protein quality rather than interface energetics [23]. Here, HGH 17 and HGH 20 combined the lowest expression levels in the panel (8.7 and 7.6 nM, against a panel median of *∼*54 nM) with anomalously low hydrodynamic radii, identifying them as expression failures rather than non-binders — a distinction a binary readout cannot make.

Full-stack pipeline automation is also what will let us learn more from each campaign. Within the pool that had passed our filters, no in silico descriptor ranked success, which we interpret as a restricted-range effect. Because every design is now expressed, quantified and measured in the same automated way, the platform already records expression level, monodispersity, unspecificity and binding kinetics for each one, and other developability-related assays could be added in the same format. Accumulated over many campaigns, such data could be used to recalibrate confidence metrics and to design for developability criteria such as pH sensitivity [33], thermal and protease stability, and immunogenicity, as well as for predicted binding, positive or negative, against one or more targets, framed as a multi-objective optimisation challenge. Additionally, eukaryotic CFPS lysates such as ALiCE offer a potential route, beyond historically *E. coli*-derived reagents, towards more complex, therapeutically relevant proteins that require disulfide bonds or N-glycosylation. In the future, newly identified binders could be reformatted into, e.g., Fc-fusion constructs for hit-to-lead development within the same CFPS platform, potentially reducing the need to revert to mammalian cell culture.

In summary, we present an in vitro validation pipeline that needs no GMO laboratory, cell culture, purification or immobilisation, and that allows simple parallelisation of e.g. three 96-well microplates every 5 days. By providing high-quality, standardised data, we hope that this pipeline begins to close the gap between how fast proteins can be designed and how fast they can be tested. In addition, the fully automated pipeline for iterative, metric-driven design and optimisation of de novo protein binders is intended to give time-constrained scientists a tool to design binders against any target of interest, meeting the in silico metric values of their will.

## Methods

### Iterative design and metric-driven optimisation of de novo protein binders

The complete pipeline for the de novo design of protein binders using an iterative metric-driven optimisation approach was fully automated. First, RFdiffusion generates binder scaffolds against each target of interest separately. Second, ProteinMPNN design sequences for those binder scaffolds. Third, AlphaFold2 Initial Guess (AF2-IG) predicts folding and evaluates the sequences, using the coordinates of the RFdiffusion-designed scaffold as the starting point for folding to speed up the process. After the first design and evaluation round, if the number of generated unique-scaffold binders passing the preset filters (ipAE and pLDDT) is lower than requested, the pipeline enters an iterative loop (scaffolds are treated as unique after each partial diffusion). It takes the best designs from the previous design round and uses RFdiffusion to partially diffuse them into new but related scaffolds; ProteinMPNN then generates new sequences for each new scaffold and AF2-IG predicts the folding and evaluates them. If the number of designs passing the filters is still lower than requested, the pipeline proceeds to the next round, until the request is met or the maximum number of iteration rounds is reached. Once either condition is satisfied, Boltz-2 predicts folding and evaluates all designs that passed the requested pLDDT and ipAE thresholds and ranks them by ipSAE min.

The pipeline was run three times: one without hot-spots (blind run) and twice with different hot-spots: His21, Phe25 and Arg167 (HS 1), and Leu6, Leu9 and Thr123 (HS 2).

### Vector preparation

The cell-free protein synthesis vector was designed using the pALiCE01 (LenioBio) as a basis and adding the fusion protein construct, which included a linker, mChartreuse, FXa cleavege site and Strep-Tag II (Fig. 3b). Custom Golden Gate handles were added to the 5’ and 3’ ends for scarless cloning. The construct was ordered as linear dsDNA from GenScript. The vector was amplified by preparative PCR at a 20 *×* 50 µL scale (1 mL total reaction volume); the master-mix composition and cycling programme are given in Supplementary Methods (Table 1 and 2). Reactions were prepared on the bench and run in a standard thermocycler. After amplification, the reactions were pooled and purified with the GeneJET PCR Purification Kit (Thermo Fisher Scientific, cat. no. K0702), according to the supplier’s instructions, with elution in a single 150 µL volume of MilliQ water and quantification by NanoDrop spectrophotometry (A260, A260/A280, A260/A230). Amplicon size was verified by 1% agarose gel electrophoresis.

### Linear DNA preparation and molecular cloning

The GenScript codon-optimisation tool was used to reverse-translate the amino-acid sequences of the binders, optimising for both *N. tabacum* and *E. coli*. Flanking sequences containing common primer-annealing sites and the recognition sites required for Golden Gate cloning [34] were added at the 5*^′^* and 3*^′^* ends. The resulting sequences were ordered as linear dsDNA gene fragments from GenScript in TE buffer.

Binder genes were amplified using Platinum SuperFi II PCR Master Mix (Thermo Fisher Scientific, cat. no. 12368050) in 30 µL reactions. Each reaction contained 15.0 µL of 2*×* Platinum SuperFi II PCR Master Mix, 1.5 µL of each 10 µM primer, 11.0 µL of MilliQ water, and 1.0 µL of template DNA. Reactions were prepared in an Armadillo 384-well PCR plate (Thermo Fisher Scientific, cat. no. AB2384) and sealed with a self-sealing PCR lid (Bio-Rad, cat. no. MSL2022) before thermal cycling. Amplification consisted of an initial denaturation at 98 °C for 30 s, followed by 30 cycles of 98 °C for 30 s, 60 °C for 30 s, and 72 °C for 30 s, with a final extension at 72 °C for 5 min.

PCR products were purified by solid-phase reversible immobilisation using MagMAX Pure Bind DNA binding beads (Thermo Fisher Scientific, cat. no. A58522). For each 30 µL PCR reaction, 54 µL of bead suspension was added and mixed for 10 min at 1000 rpm. The beads were magnetically separated for 2 min and the supernatant was removed. The beads were washed twice with 50 µL of 70% (v/v) ethanol, followed by a 2-min drying period. DNA was eluted in 15 µL of MilliQ water by shaking for 10 min at 2000 rpm.

Purified binder genes were inserted into the mChartreuse-containing [35] expression vector by Golden Gate assembly (Avelar-Molina et. al, in preparation). Per reaction, the Golden Gate master mix contained 1.50 µL of 10*×* T4 DNA Ligase Buffer (New England Biolabs, cat. no. B0202), 1.50 µL of BsaI-HFv2 (New England Biolabs, cat. no. R3733), 0.10 µL of T4 DNA Ligase (New England Biolabs, cat. no. M0202T), and 5.16 µL of mChartreuse vector at 348.9 ng/µL, corresponding to 1,800 ng of vector. The resulting 8.26 µL master mix was combined with 7.80 µL of purified binder gene, giving a total reaction volume of 16.06 µL. As a fixed volume of purified gene was used rather than normalisation by DNA mass, the insert-to-vector molar ratio varied between constructs. Reactions were prepared in the Armadillo 384-well PCR plate and sealed with a self-sealing PCR lid (Bio-Rad, cat. no. MSL2022) before thermal cycling. Golden Gate assembly was performed for 30 cycles of 37 °C for 60 s and 16 °C for 60 s, followed by 60 °C for 10 min and a hold at 10 °C. The assembled DNA was then purified by the SPRI procedure described above, using 27 µL of MagMAX Pure Bind DNA binding beads per reaction, and eluted in 15 µL of MilliQ water. DNA products were verified by agarose gel electrophoresis as described above.

### Agarose gel electrophoresis

Vector, binder gene fragments and Golden Gate assembly identities were verified by agarose gel electrophoresis. Gels were cast as 1% (w/v) agarose in 1*×* TAE buffer supplemented with SYBR Safe DNA gel stain (1*×* final concentration). Samples (3 µL) were mixed with 0.5 µL of 6*×* dna loading dye, and 3 µL was loaded per well; 3 µL of GeneRuler 1 kb Plus dna Ladder (Thermo Fisher Scientific) was loaded in flanking lanes. Electrophoresis was performed in 1*×* TAE buffer at 110 V for 50 min and gels were visualised on a Bio-Rad ChemiDoc MP imaging system.

### Binder cell-free protein synthesis

Binders were expressed using 18 µL of ALiCE cell-free protein synthesis (CFPS) reagent (LenioBio) [28, 36] and 2 µL of the purified DNA (72-120 ng/µL) in a low-protein-binding 384-well plate (Thermo Fisher Scientific, cat. no. 267461), sealed with a permeable polyolefin lid (HJ-BIOANALYTIC GmBH 900371) for oxygen transfer and incubated at 25 °C, 800 rpm and 85% relative humidity in an Infors HT Multitron incubator (Infors AG) for 36 h.

### In-solution binder screening directly in CFPS material

After the 36 h incubation, 1 µL each of the CFPS reaction products were transferred to 99 µL of 10 mM sodium phosphate, pH 8.5, with 0.05% Pluronic acid (1:100 dilution) in a FIDABIO 96-well plate (Pr. No: 220-002). Flow-induced dispersion analysis (FIDA) was then performed using a FIDA Neo (Fida Biosystems ApS) with LED-induced fluorescence detection (excitation 480 nm, emission ¿515 nm). We used permanently coated capillaries to reduce adsorption (75 µm inner diameter, 375 µm outer diameter, 1 m total length, and 84 cm to the detection window) and 0.1% phosphoric acid for washing. The used capillary mixing (CapMix) method consisted of 5 automated steps: (1) 30 s 0.1% phosphoric acid wash at 3500 mbar, (2) 20 s pre-equilibration at 3500 mbar with our buffer, (3) 30 s analyte injection at 3500 mbar, (4) 20 s indicator injection at 50 mbar, and (5) 180 s analyte injection at 400 mbar.

Each binder (the fluorescent indicator due to mChartreuse fusion) was run twice: first, with buffer (10mM sodium phosphate, pH 8.5, 0.05% Pluronic acid) as analyte to obtain the hydrodynamic radius (Rh) of the binder alone (baseline), and then with the target (5 µM hGH) as analyte. The hGH was obtained in a lyophilised form from Thermo Fisher Scientific (cat. no. 200-02-250UG), initially reconstituted in water to 0.1-1.0 mg/ml and further diluted in 10 mM sodium phosphate, pH 8.5. Results were analysed with FIDABIO proprietary software.

### Binding affinity and kinetics

Equilibrium binding curves were obtained using FIDA in a PreMix mode, in which analyte and indicator were mixed in advance in the 96-well plate. The indicator concentration was kept fixed while the analyte (hGH) was titrated over 8 concentration points ranging from 10 µM to 0.61nM plus a zero point. To avoid having an overly steep binding curve, the binders were diluted as much as possible while keeping the signal-to-noise ratio higher than 100 for accurate Rh measurements. The method consisted of 4 automated steps: (1) 30 s 0.1% phosphoric acid wash at 3500 mbar, (2) 50 s pre-equilibration at 3500 mbar with our buffer, (3) 20 s pre-mixed (indicator + analyte) injection at 50 mbar, and (5) 180 s analyte injection at 400 mbar.

Non-equilibrium kinetic analysis was performed using the FIDA CapMix mode with the same hGH concentrations as in the equilibrium binding curves. This method consisted in five automated steps: (1) 30 s of 0.1% phosphoric acid wash at 3,500 mbar, (2) 50 s of pre-equilibration at 3,500 mbar with our buffer, (3) 20 s of analyte injection at 3500 mbar, (4) 10 s of indicator injection at 50 mbar, and (5) 60 s of analyte injection at 1,200 mbar. Results were analysed with FIDABIO proprietary software.

Binder concentrations for each experiment were derived from a calibration curve. Binder HGH 3 was purified using MagStrep® Strep-Tactin®XT beads magnetic beads (IBA Lifesciences cat. no. 2-5090-010) following the supplier’s protocol, and its concentration was determined using Nanodrop and measuring fluorescence at 280 and 510 nm, together with the Extincion coefficients of mChartreuse at 510 nm (71,000 M^1^cm^1^) and the theoretical extinction coefficient of the HGH 3-mChartreuse complex at 280 nm (22,015 M^1^cm^1^ ProtParam Expasy [37]). Eight binder concentrations were analysed by FIDA Neo using the same method as for baselines in the binder screening and the fluorescence AUC was plotted against the binder concentration to construct a calibration curve for binder quantification. HGH 10, HGH 12, HGH 18 and HGH 30 were also purified the same way and, together with HGH 3, run in an SDS-PAGE gel to confirm their presence.

### Automated workflow from gene fragment to circular DNA

The DNA preparation and cloning workflow described above was implemented end-to-end on a Tecan Fluent 780 liquid-handling platform operated with FluentControl (build 3.7.19.207) on Windows 11 Enterprise IoT. The system comprised an eight-channel flexible channel arm (FCA), a 384-channel multi-channel arm (MCA) equipped with a 96-channel adapter, and a robotic gripper arm (RGA). An INHECO ODTC^®^ 384XL thermal cycler was mounted directly on the Dynamic Deck^TM^ and integrated into FluentControl using the manufacturer’s OEM driver. A BioShake 3000 T-ELM shaker (QINSTRUMENTS GmbH) was integrated through DriverFramework driver over RS232 and an Alpaqua Magnum FLX^®^ magnet was placed on the worktable for magnetic-bead purification. Custom liquid classes were used for bead suspensions, ethanol handling with the MCA, and low-volume DNA transfers.

PCR and Golden Gate master mixes were prepared in bulk before initiation of the automated method and loaded together with the magnetic beads and MilliQ water in a 96-well 2-mL deep-well source plate. Samples were organised in an Armadillo 384-well PCR plate (Thermo Fisher Scientific, cat. no. AB2384), with four wells assigned to each candidate construct. Each sample followed the sequence unpurified PCR product *→* purified PCR product *→* Golden Gate reaction *→* purified Golden Gate product; for the first construct this corresponded to A1 *→* A2 *→* B1 *→* B2. This four-well arrangement was repeated across the plate, enabling up to 96 constructs to be processed in parallel while maintaining a fixed sample position at each stage.

For PCR setup, the FCA dispensed 29 µL of PCR master mix into the first designated well and the MCA added 1 µL of template DNA. A self-sealing PCR lid (Bio-Rad, cat. no. MSL2022) was placed on the Armadillo plate by the RGA before it was transferred to the integrated ODTC for thermal cycling. Following PCR, the products were transferred by the MCA to a 96-well purification plate containing 27 *µL* beads dispensed by the FCA. The RGA subsequently moved the plate between the BioShake and Alpaqua magnet for binding, washing, and elution, after which the purified DNA was returned by the MCA to the second designated position in the 384-well plate.

The FCA then dispensed Golden Gate master mix into the third designated position and the MCA transferred the corresponding purified gene product. A self-sealing PCR lid (Bio-Rad, cat. no. MSL2022) was applied before the RGA transferred the sealed plate to the integrated ODTC for Golden Gate cycling. Following assembly, the products underwent a second automated magnetic-bead purification using the same sequence of shaker and magnet operations. The final purified constructs were transferred to the fourth designated position in the 384-well plate, from which they were used for downstream cell-free protein synthesis.

## SUPPLEMENTARY

The production Golden Gate reaction composition and cycling programme are given in Tables 1 and 2.

**Table 1.** Optimised PCR programme for backbone amplification, applied to mChartreuse backbone constructs.

| | Temperature ($^{\circ}$C) | Time (s) | Cycles |
| --- | --- | --- | --- |
| Initial denaturation | 98 | 30 | 1 |
| Denaturation | 98 | 15 | 34 |
| Annealing | 56 | 20 | 34 |
| Extension | 72 | 45 | 34 |
| Final extension | 72 | 300 | 1 |
| Hold | 4 | ∞ | 1 |

**Table 2.** Composition of the backbone PCR master mix for a single 50 µL reaction. Volumes scale linearly to the desired number of reactions; routine production used 20 reactions pooled to 1 mL per backbone construct.

| Component | Volume (μL) |
| --- | --- |
| 2× Platinum SuperFi II PCR MasterMix | 25.0 |
| CFPS_fwd primer (10 μM) | 2.5 |
| CFPS_rev primer (10 μM) | 2.5 |
| Linear vector template (∼1 ng/μL) | 1.0 |
| MilliQ water | 19.0 |
| Total | 50.0 |

## Declarations

### Funding/Acknowledgments

DRE and GAM acknowledge support from Novo Nordisk A/S as part of the DTU Novo Nordisk Closed-Loop Optimization initiative.

### Author contribution

**Conceptualization** ERdT, TPJ, AH, DRE, GAM. **Methodology:** DRE, GAM, OSB, AU, KSR, CW. **Investigation:** DRE, GAM, OSB, JH, AU, LWG, KBL. **Formal analysis:** DRE, OSB, JH, KSR, CW. **Supervision:** ERdT, GAM, AU, DRE. **Funding acquisition:** TPJ, AH. **Writing (original draft):** DRE, ERdT, OSB. **Writing (review & editing):** All authors read and approved the final version of the manuscript.

## References

[1] Watson, J.L., et al.: De novo design of protein structure and function with RFdiffusion. Nature 620, 1089–1100 (2023)

[2] Dauparas, J., et al.: Robust deep learning-based protein sequence design using ProteinMPNN. Science 378, 49–56 (2022)

[3] Bennett, N.R., et al.: Improving de novo protein binder design with deep learning. Nat. Commun. 14, 2625 (2023)

[4] Zambaldi, V., et al.: De novo design of high-affinity protein binders with AlphaProteo. arXiv (2024). arXiv:2409.08022

[5] Pacesa, M., et al.: One-shot design of functional protein binders with BindCraft. Nature 646, 483–492 (2025)

[6] Cao, L., et al.: De novo design of picomolar SARS-CoV-2 miniprotein inhibitors. Science 370, 426–431 (2020)

[7] Jumper, J., et al.: Highly accurate protein structure prediction with AlphaFold. Nature 596, 583–589 (2021)

[8] Abramson, J., et al.: Accurate structure prediction of biomolecular interactions with AlphaFold 3. Nature 630, 493–500 (2024)

[9] Bryant, P., et al.: Improved prediction of protein-protein interactions using AlphaFold2. Nat. Commun. 13, 1265 (2022)

[10] Zhu, W., et al.: Evaluation of AlphaFold-Multimer prediction on multi-chain protein complexes. Bioinformatics 39, 424 (2023)

[11] Dunbrack, R.L. Jr: Re̅s ipSAE loquuntur: what’s wrong with AlphaFold’s ipTM score and how to fix it. bioRxiv (2025). doi:10.1101/2025.02.10.637595

[12] Passaro, S., Corso, G., Wohlwend, J., et al.: Boltz-2: towards accurate and efficient binding affinity prediction. bioRxiv (2025). doi:10.1101/2025.06.14.659707

[13] Overath, M.D., et al.: Predicting experimental success in de novo binder design: a meta-analysis of 3,766 experimentally characterised binders. bioRxiv (2025). doi:10.1101/2025.08.14.670059

[14] Authors to be added: Crowdsourced protein design: lessons from the Adaptyv EGFR binder competition. bioRxiv (2025). doi:10.1101/2025.04.17.648362

[15] Tsuboyama, K., et al.: Mega-scale experimental analysis of protein folding stability in biology and design. Nature 620, 434–444 (2023)

[16] Rocklin, G.J., et al.: Global analysis of protein folding using massively parallel design, synthesis, and testing. Science 357, 168–175 (2017)

[17] Chevalier, A., et al.: Massively parallel de novo protein design for targeted therapeutics. Nature 550, 74–79 (2017)

[18] Cao, L., et al.: Design of protein-binding proteins from the target structure alone. Nature 605, 551–560 (2022)

[19] Yao, Z., Metts, M., Huber, A.K., Li, J., Kinjo, T., Dieckhaus, H., Nallathambi, A., Bowers, A.A., Kuhlman, B.: Enhancing machine learning-based binder design with high-throughput screening: a comparison of mRNA and yeast display technologies. Protein Sci. (2026). doi:10.1002/pro.70696

[20] Hurley, J.D., Kruse, A.C.: Profiling protein-peptide interactions by yeast surface display. Curr. Protoc. 6, 70436 (2026). doi:10.1002/cpz1.70436

[21] Reusch, J., Andersen, J.T., Rant, U., Schlothauer, T.: Insight into the avidity–affinity relationship of the bivalent, pH-dependent interaction between IgG and FcRn. mAbs 16(1), 2361585 (2024) 10.1080/19420862.2024.2361585

[22] Liu, H., Chen, P., Zhai, X., Huo, K.-G., Zhou, S., Han, L., Fan, G.: PPB-Affinity: Protein-protein binding affinity dataset for AI-based protein drug discovery. Scientific Data 11(1), 1316 (2024) 10.1038/s41597-024-03997-4

[23] Kanter, J.K., Smorodina, E., Minnegalieva, A., Arts, M., Blaabjerg, L.M., Frolenkova, M., Rawat, P., Wolfram, L., Britze, H., Wilke, Y., Weissenborn, L., Lindenburg, L., Engelhart, E., McGowan, K.L., Emerson, R., Lopez, R., Bemmel, J.G., Demharter, S., Spreafico, R., Greiff, V.: Effects of protein interface mutations on protein quality and affinity. bioRxiv (2026) 10.64898/2026.03.24.713863. Preprint

[24] Jarmoskaite, I., AlSadhan, I., Vaidyanathan, P.P., Herschlag, D.: How to measure and evaluate binding affinities. eLife 9, 57264 (2020) 10.7554/eLife.57264

[25] Capin, J., Mayonove, P., DeVisch, A., Becher, A., Ngo, G., Courbet, A., Ragotte, R.J., Cohen Gonsaud, M., Espeut, J., Bonnet, J.: CF2H: a cell-free two-hybrid platform for rapid protein binder screening. Nat. Commun. 17, 3724 (2026)

[26] Jensen, H., Østergaard, J.: Flow induced dispersion analysis quantifies noncovalent interactions in nanoliter samples. J. Am. Chem. Soc. 132(12), 4070–4071 (2010). doi:10.1021/ja100484d

[27] Pinheiro, F., Nowak, J.S., Zueva, E., Pheasant, E.C., Grene, I.K., Lampinen, V., Kjaergaard, M.: Screening de novo designed protein binders in unpurified lysate using flow induced dispersion analysis. Protein Sci. 34, 70286 (2025)

[28] Das Gupta, M., Flaskamp, Y., Roentgen, R., Juergens, H., Armero-Gimenez, J., Albrecht, F., Hemmerich, J., Arfi, Z.A., Neuser, J., Spiegel, H., Schillberg, S., Yeliseev, A., Song, L., Qiu, J., Williams, C., Finnern, R.: Scaling eukaryotic cell-free protein synthesis achieved with the versatile and high-yielding tobacco BY-2 cell lysate. Biotechnol. Bioeng. 120(10), 2890–2906 (2023). doi:10.1002/bit.28461

[29] Cunningham, B.C., Wells, J.A.: High-resolution epitope mapping of hGH-receptor interactions by alanine-scanning mutagenesis. Science 244, 1081–1085 (1989)

[30] Vos, A.M., Ultsch, M., Kossiakoff, A.A.: Human growth hormone and extracellular domain of its receptor: crystal structure of the complex. Science 255, 306–312 (1992)

[31] Trainer, P.J., Drake, W.M., Katznelson, L., Freda, P.U., Herman-Bonert, V., Lely, A.J., Dimaraki, E.V., Stewart, P.M., Friend, K.E., Vance, M.L., Besser, G.M., Scarlett, J.A.: Treatment of acromegaly with the growth hormone-receptor antagonist pegvisomant. N. Engl. J. Med. 342(16), 1171–1177 (2000)

[32] Benjamini, Y., Hochberg, Y.: Controlling the false discovery rate: A practical and powerful approach to multiple testing. J. R. Stat. Soc. Series B 57(1), 289–300 (1995). doi:10.1111/j.2517-6161.1995.tb02031.x

[33] Jacobsen, C.P., Deweid, L., Birnbaum, F., Gormsen, L.W., Fryer, T., Jagd, U., Hald, A., Rasch, M.G., Jenkins, T., Ovchinnikov, S.: pH-sensitive binder design with Proton-PottsMPNN. bioRxiv (2026). doi:10.64898/2026.09.30.755438

[34] Engler, C., Kandzia, R., Marillonnet, S.: A one pot, one step, precision cloning method with high throughput capability. PLoS ONE 3(11), 3647 (2008). doi:10.1371/journal.pone.0003647

[35] Fraikin, N., Couturier, A., Lesterlin, C.: A palette of bright and photostable monomeric fluorescent proteins for bacterial time-lapse imaging. Sci. Adv. 11(16), 6201 (2025). doi:10.1126/sciadv.ads6201

[36] Armero-Gimenez, J., Schleusner, J., Wilbers, R.H.P., Schots, A., Spiga, L., Tregoning, J.S., Brown, J.C., Zhou, J., Juergens, H., Finnern, R., Williams, C.: Noncanonical amino acids enable plug and play vaccine platform in the ALiCE cell free system. npj Vaccines 11(1), 80 (2026). doi:10.1038/s41541-026-01435-7

[37] Gasteiger, E., Hoogland, C., Gattiker, A., Duvaud, S., Wilkins, M.R., Appel, R.D., Bairoch, A.: Protein identification and analysis tools on the ExPASy server. In: Walker, J.M. (ed.) The Proteomics Protocols Handbook, pp. 571–607. Humana Press, ??? (2005)

